# Quantitative analysis of cellular morphology during *in vitro* decidualization

**DOI:** 10.1101/2022.10.27.513741

**Authors:** Luciana Ant, François Le Dily, Miguel Beato, Patricia Saragüeta

## Abstract

Decidualization is a differentiation process involving shape reorganization from a fibroblast to an epithelioid-like appearance of endometrial stromal cells. Specificities of these cells impede the use of existing automated tools to follow morphological changes during differentiation; we therefore developed a simple but accurate methodology to quantify the phenotypical changes that occur in an *in vitro* decidualization system.

The approach consists of the analysis of the circularity of the cells directly from light microscopy images. Here, we used this methodology to follow the effects of progesterone or progestin R5020 in combination with estradiol (E2) and cAMP on inducing the decidualization of human endometrial cells. We further implemented a statistical model to detect the differences in the kinetics of decidualization of the two hormonal stimuli before all the cell population acquired the decidual phenotype. We found that 2 days after stimulation are sufficient to detect statistical differences in morphology between decidualization induced and control cells. Here, we detail the model and scripts in order to provide a useful, practical and low cost tool to evaluate morphological aspects of endometrial stromal differentiation.

**Availability and implementation:** See supplementary methods

**Supplementary information:** Supplementary data is available online.

## Introduction

Decidualization is the transdifferentiation process of stromal endometrial cells into decidual cells, which involves changes in cytoplasm and nuclei morphology (2,3) and changes in gene expression associated to a secretory signature to support the embryo in case of implantation success (6). This process can be recapitulated *in vitro* by the use of hormonal treatments. Although decidualization markers can be used to analyze the differentiation by immune-fluorescence (3,8), for example at the end point of the process once the whole population of cells are committed, these approaches may not be always suitable to follow the dynamic of those changes in a time course manner. In order to better approach this problem and analyze the proportion of decidualized cells during the time course of hormonal stimulation, we aimed at quantifying the morphological changes that occur during this process. Although decidualization associated phenotypic changes are recapitulated *in vitro* and detectable on light microscopy images, the extremely flat phenotype of endometrial cells in 2D culture conditions impede the use of existing tools developed to follow morphological changes in other cell models (7). In this work, we therefore implemented a bioinformatic and statistical model to quantify the morphological changes accompanying differentiation in an *in vitro* decidualization system directly from light microscopy images. The approach presented here covers the necessity to quantify morphology changes in a simple, low cost, and accurate way allowing to follow the dynamic of differentiation before all the cell population acquires a decidual phenotype.

## Results

In order to determine potential differences in the kinetic of decidualization induced by different hormonal combinations, we treated human endometrial T-HESC cells (4) up to 6 days with EtOH vehicle as control or with E2 and cAMP in combination with different progestins, either P4 or R5020 (see materials and methods and Fig. 1a). The main workflow of the process is described in figure 1b.

**Fig.1.**
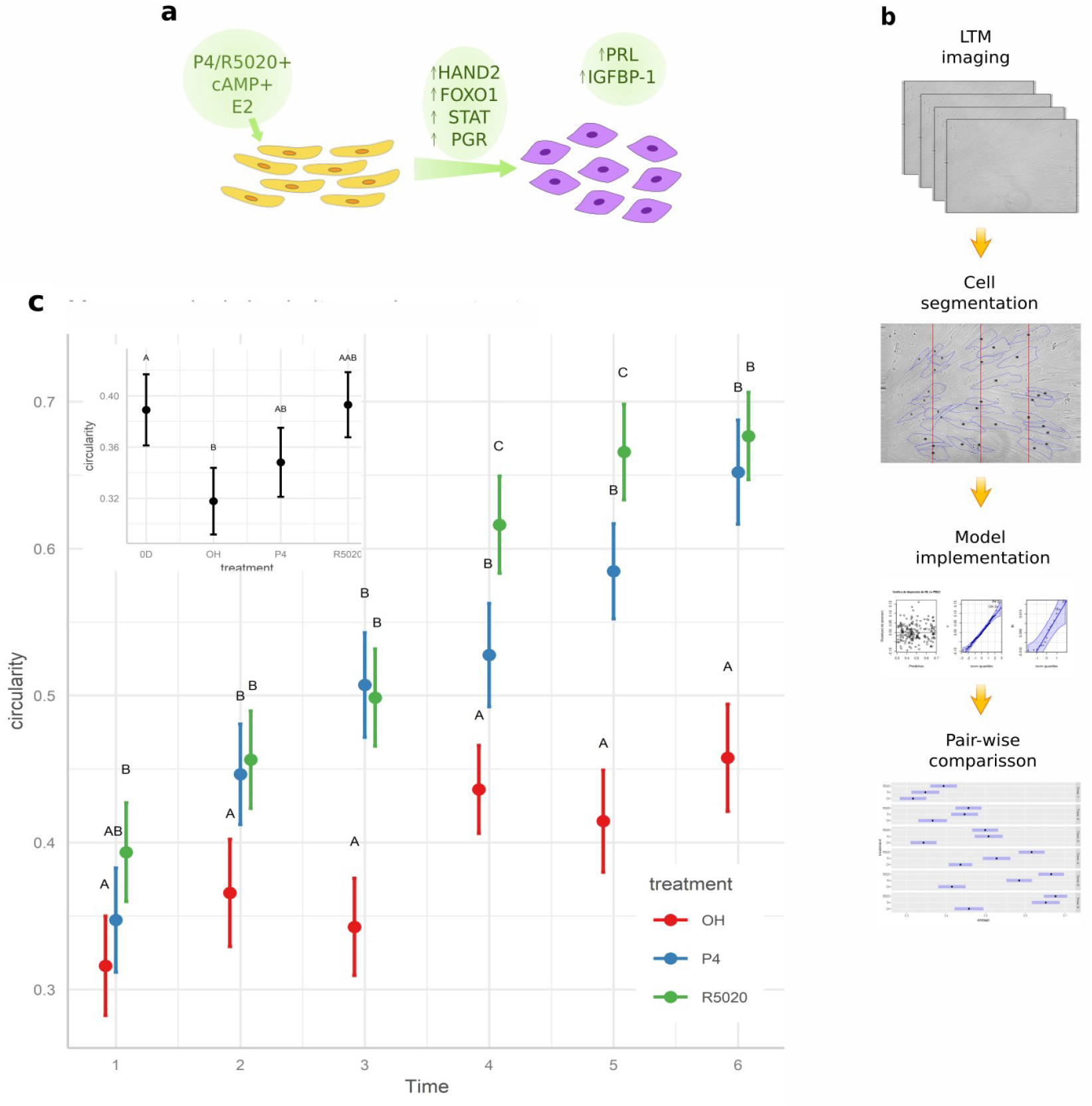
**a.** Morphology quantification of in vitro decidualization. A) Schematic overview of the decidualization system. Decidualizing stimuli (cAMP, E2 and P4/R5020), key transcription factors (HAND2, STATs, PGR, FOXO1) and decidualization markers (PRL and IGBP1) upregulated during this process are noted. B) Workflow used in this study. C) Estimated marginal circularity means for each treatment using the conditional model m1. Different letters indicate statistical differences between treatments at a given day. Inset denotes circularity means estimated by the simple model comparing T0 and Day 1. Each dot represents the mean circularity predicted for each treatment. Bar represents the interval of confidence. Different letters indicate statistical differences between different treatments and days.

The decidualized final state at 6 days was confirmed by qualitative changes in morphology of treated cells compared to cells cultured in presence of the vehicle (Fig S1a) as well as by the detection of decidualization markers, PRL and IGFBP1 by qRT-PCR (Fig S1b).

To quantify the kinetic of the changes in cell morphology, light transmitted images were taken before treatment (T0) and every day during 6 days for each condition of treatment (5 plates by condition x 2 areas per plate). The circularity of the cells measured from manually curated images was used as a proxy of morphological changes. Exploratory analysis comparing the distributions of the cell circularity measures showed normal distribution of this parameter (Fig S2). Thus, mean circularity weighted by the number of cells per image was used as the response variable that did adjust to a normal distribution (Fig S3). In order to compare the effects of time and treatment, we used these data to implement simple Mixed-Model Regression. Since the T0 plates did not share time or treatment with any other conditions, we applied two different statistical models for the comparison: one to compare the circularity between T0 and Day1 and one to compare the circularity of the cells from day 1 to day 6, using the variables “time” and “treatment” (see materials and Methods). The first comparison did not show any significant differences between the T0 condition and the decidualizing treatments after one day (inset of Fig 1C), In contrast, the dynamics of morphological changes from day 1 to 6 appears to happen in a two-phase manner (Fig. 1C). From day 2, both decidualization treatments differentiate from control cells treated with EtOH, showing that our approach is sufficient to detect early morphological changes in the cell population. This difference is maintained throughout the remaining days. At day 4 the cells from all treatments show a leap in their circularity. This could be due to an increment in cell density, that causes the cells to be smaller and less elongated.

From day 4, cells treated with R5020 showed increased circularity as compared to cells induced with P4. The final state at day 6 shows the same circularity for both hormonal treatments.

The changes in circularity observed in decidualized cells are a result of mild changes in circularity in the majority of the cells, and not an increase in a few cells, as can be seen in the density plots (figure S2a).

## Discussion

Here a model to quantify in vitro decidualization was implemented without defining the gene expression signature. This method was practical and reports a result of multiple processes that comprise the phenotypic transformation during decidualization. This approach allows to verify the establishment of the decidual phenotype without the need to use cell culture in RT-PCR or immunohistochemistry analysis.

The quantitative analysis of the morphological changes during decidualization allowed us to gain insight that would not be possible to obtain through a qualitative analysis. The difference in circularity observed for the decidualization treatments at day 4 and 5 could be attributable to the difference in metabolization of the two progestins. R5020 is a synthetic progestin that cannot be metabolized in the cell, making the concentration levels of this drug stable during the experiment. Progestin on the other hand is metabolized as the days go by, decreasing its concentration levels. This could have slowed down the morphological changes observed in progesterone treated cells compared to R5020.

## Materials and Methods

### Implementation

Circularity measures were acquired in ImageJ using custom macros, in order to render the process as efficient as possible. Data analysis was done entirely in R using basic packages such as nlme and emmeans. All the scripts and macros were developed for this analysis and can be found in the supplementary material.

### Cell culture and treatment

Human endometrial stromal cells (t-HESC, ATCC CRL-4003) were cultured for 48 hs. in DMEM F12 (D2906, Sigma Aldrich) with 10% DCC-FBS (5). The cells were then trypsinized and re-cultured in 15 100mm plates with a density of 1×10^6^ cells per plate. After 24 hours the medium was replaced with DMEM-F12 2%-DCC-FBS and either Progesterone 10^-6^ M or R5020 1×10^8^, Estradiol 10nM and 8-Bromo-AMPc 0,5mM, while the control plates were given DMEM-F12 2%-DCC-FBS EtOH 0,1.

### RT-qPCR

Cell extracts were collected in denaturing solution (4 M Guanidine thiocyanate, 25 mM Sodium citrate pH 7, 0.1 M 2-Mercaptoethanol, 0.5% Sarkosyl) and total RNA was prepared following phenol-chloroform protocol (1). Integrity-checked RNA was used to synthesize cDNA with oligodT (Biodynamics) and MMLV reverse transcriptase (Thermo Fisher Scientific).

Quantification of candidate gene products was assessed by real-time PCR. Expression values were corrected by GAPDH and expressed as mRNA levels over time zero (T0). Primer sequences are:

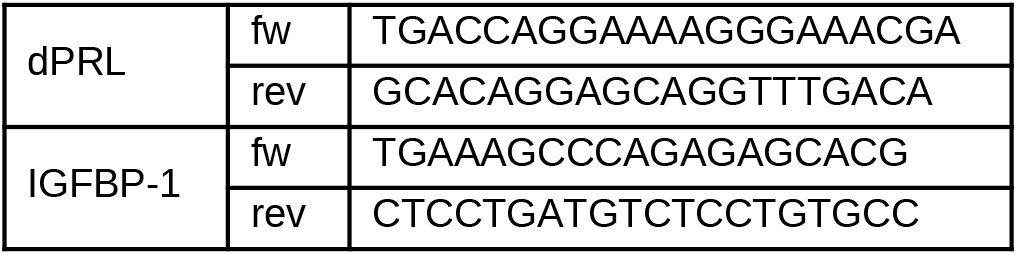

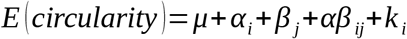

### Image acquisition

Images of two random areas of each plate were taken before treatment, and once a day from day 1 to 6. Images were obtained using a Leica DMI6000 B inverted microscope.

### Cell segmentation and manual image curation

As cultured tHESC cells are too flat to be recognized by an AI algorithm the images were manually curated. Circularity was measured through ImageJ. The segmenting of the cells from each image was performed by hand. Then, circularity was measured as 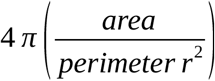, where a circularity value of 1.0 indicates a perfect circle, and an increasingly elongated polygon as the value approaches 0.0.

### Statistical Model

Mean circularity per photo was used as the response variable that did adjust to a normal distribution (Fig S3). The number of cells in each photo were taken into account as the “weight” for each mean. Due to the fact that the plates used for T0 images did not share time or treatment with any other plate we applied two statistical models: one to compare the circularity of the cells from day 1 to day 6, using the variables “time” and “treatment”, and another one to compare the circularity between T0 and Day1. The latter only comprised one variable, called treatment, which categories were: “T0”, “Day1 EtOH”, “Day1 R5020” and “Day1 P4”. The generalized linear mixed model used to compare plates from day 1 to 6 followed the equation below:

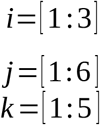

In this equation α represents the treatment effect, β the day effect, and αβ the interaction effect between them. B denotes the random effect given by the plate.

The Anova test showed a significant p-value (p-value < 0.05) for time, treatment, and the interaction between them, stating that both had an effect over the circularity of the cell, and that effect could change depending on their combination (Table S1).

On the other hand, the model comparing T0 and Day 1 matched the equation below:

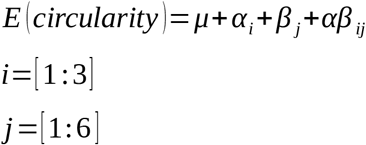

The Anova test also showed the treatment has a significant effect over circularity (Table S2).

## Acknowledgements

We are grateful to members of Saragüeta laboratories for help and suggestions and Dr. Gerardo Rubén Cueto from FCEN-UBA.

