## Supplementary Materials and Methods for "Quantitative analysis of cellular morphology during *in vitro* decidualization"

The specific methods followed in this work can be found thoroughly explained in the corresponding scripts

**Supplementary Figures.**

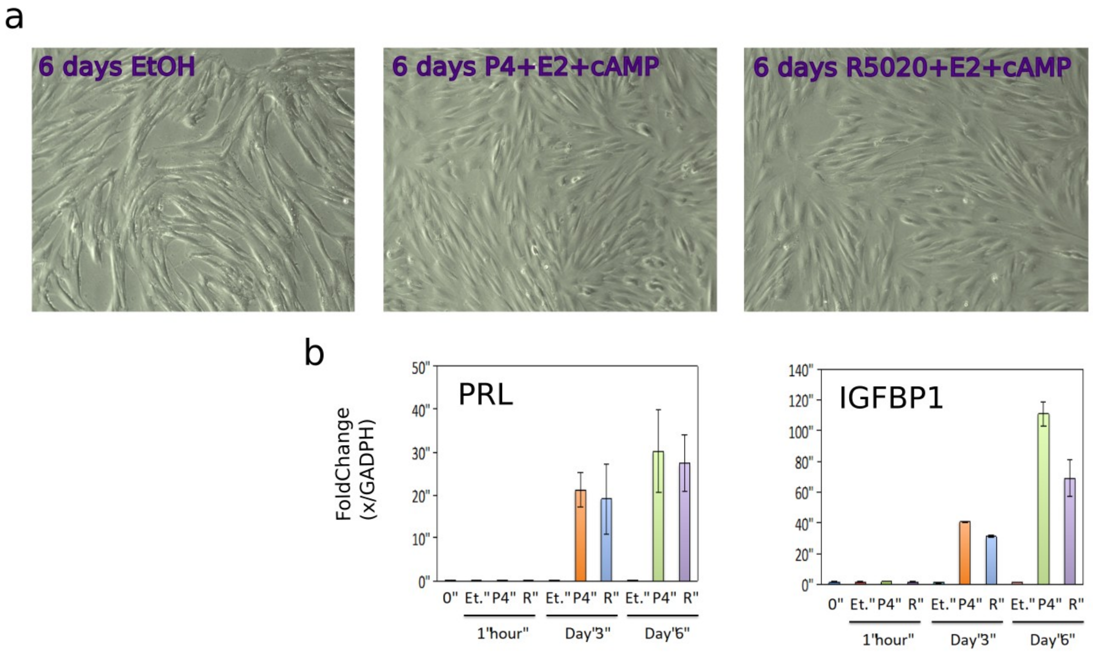


Fig. S1: a. Representative TLM images of the THESCs at six days after treatment with OH, P4+E2+cAMP and R5020+E2+cAMP. b. Expression of decidualization markers PRL and IGFBP1 at T0, 1h and 3 and 6 days after treatment.

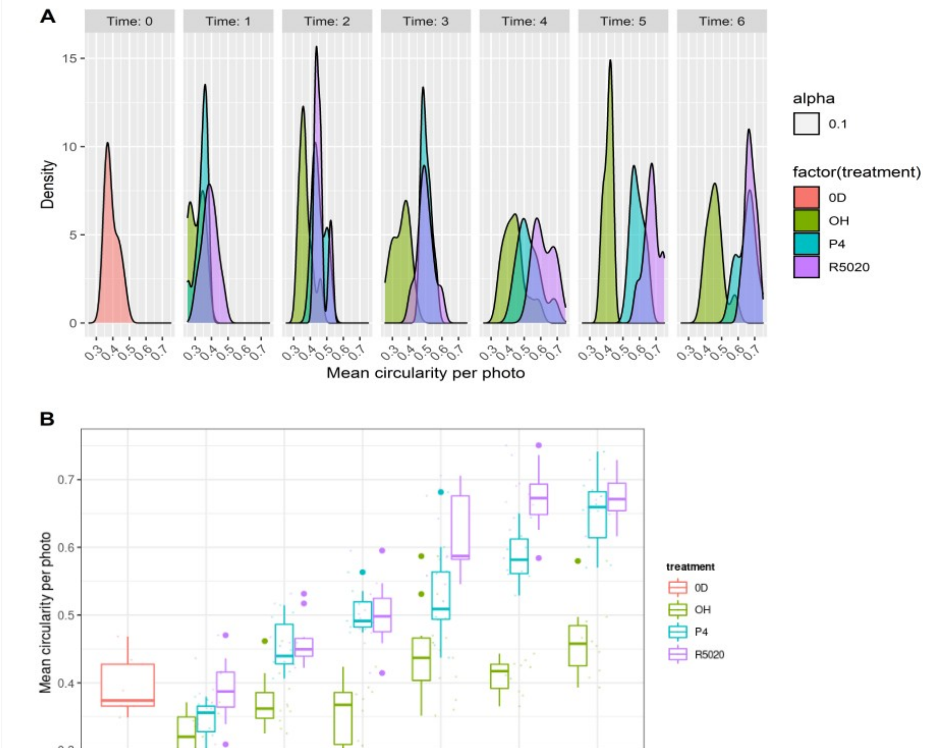


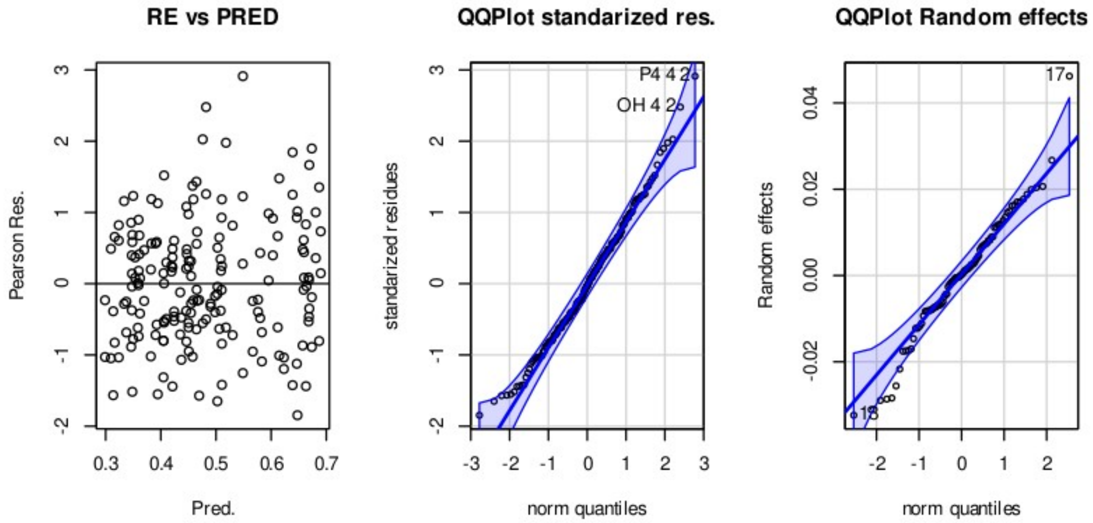


F
c
r

*I*
*c*
*c*

| | numDF | denDF | F-value | p-value |
|---|---|---|---|---|
| (Intercept) | 1 | 94 | 14717.905 | <.0001 |
| treatment | 2 | 71 | 149.506 | <.0001 |
| time | 5 | 71 | 86.893 | <.0001 |
| treatment:time | 10 | 71 | 5.666 | <.0001 |

T
s

| | numDF | denDF | F-value | p-value |
|---|---|---|---|---|
| (Intercept) | 1 | 21 | 3323.645 | <.0001 |
| treatment | 3 | 16 | 8.343 | 0.0014 |

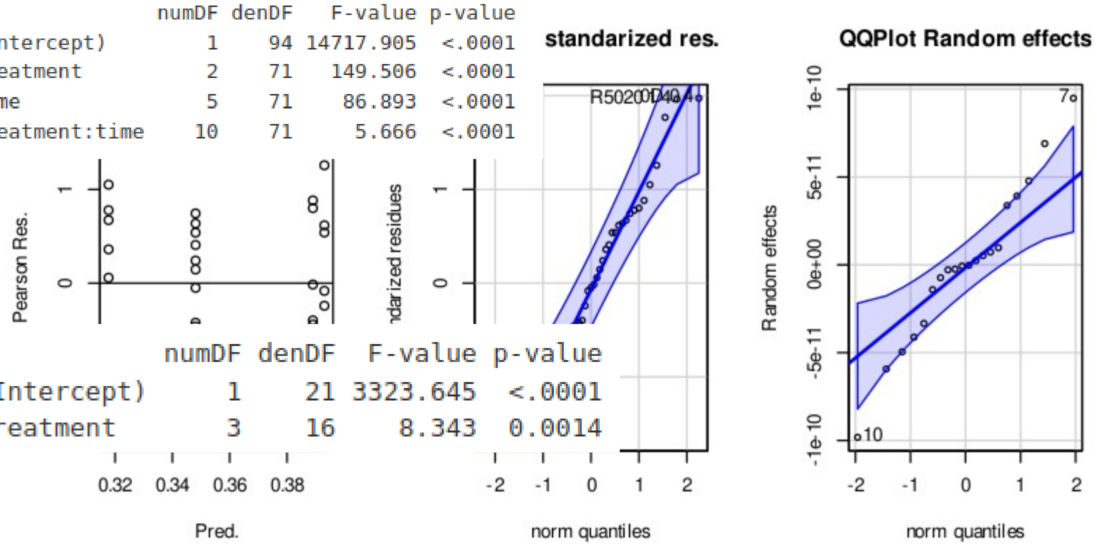

*Table S2: Anova for the simple model comparing T0 and Day1. A p-value < 0.01 is considered significant.*
